# Novel recombinant bovine papular stomatitis virus expressing foot and mouth disease virus-like particles elicits protective levels of neutralizing antibodies in cattle

**DOI:** 10.64898/2026.06.01.729332

**Authors:** Herman M. Chambaro, Gustavo Delhon, María Cruz Miraglia, Sabrina Galdo-Novo, Ana Taffarel, Byung-Joon Seung, Sushil Khatiwada, Daniel Mariano Pérez-Filgueira, Daniel L. Rock

**Affiliations:** Department of Pathobiology, College of Veterinary Medicine, University of Illinois at Urbana-Champaign, Urbana, IL, USA; School of Veterinary Medicine and Biomedical Sciences, and Nebraska Center for Virology, University of Nebraska-Lincoln, Lincoln, NE, USA; Instituto de Virología e Innovaciones Tecnológicas, CICVyA, INTA-CONICET, Buenos Aires, Argentina; WOAH FMD Reference Laboratory, SENASA, Buenos Aires, Argentina

**Keywords:** Foot and Mouth Disease, FMDV, Vaccine, Virus-like particles, Recombinant Bovine papular stomatitis virus, Cattle immunization

## Abstract

Foot-and-mouth disease (FMD) remains a major burden in endemic regions, where inactivated vaccines are constrained by cost, short duration of immunity, cold-chain dependence, and high-containment manufacturing. Here, we engineered a recombinant bovine papular stomatitis virus (rBPSV) expressing the FMDV A24 Cruzeiro capsid precursor P1-2A together with an attenuated 3C protease (3C^pro L127P^). Infection of ovine fetal turbinate (OFTu) cells resulted in robust capsid protein expression and assembly of abundant 25-30 nm icosahedral virus-like particles (VLPs). Intramuscular immunization of two BPSV-seropositive calves with rBPSV (10^7^ TCID_50_ on days 0, 21 and 35) induced strong anti-FMDV humoral responses. By day 28, liquid-phase blocking ELISA (LPBE) titers exceeded laboratory defined protective threshold (≥1.9 log_10_), and homologous neutralizing titers against A24 Cruzeiro surpassed the threshold associated with protection (≥1.36 log_10_). Intra-serotypic cross-neutralization was observed against A/Argentina/2001, whereas no neutralization was detected against heterologous serotype O_1_ Campos. Pre-existing anti-BPSV antibodies did not prevent induction of neutralizing responses. These findings establish first proof-of-concept that BPSV can serve as a cattle-adapted vector platform for delivery of FMDV VLPs and other heterologous antigens.

## Introduction

Foot-and-mouth disease (FMD) is a transboundary animal disease that continues to impose major constraints on global livestock production and international trade [1, 2]. The causative agent, FMD virus (FMDV; genus *Aphthovirus*, family *Picornaviridae*) is a small, non-enveloped, positive-sense RNA virus with an ∼8.5 kb genome that is translated into a single polyprotein, which is proteolytically processed into structural and non-structural proteins [3]. The viral capsid comprises 60 copies each of VP1, VP2, VP3 and VP4, produced from the P1-2A precursor by the viral 3C protease (3C^pro^), and assembled via protomer and pentamer intermediates into icosahedral particles displaying conformational neutralizing epitopes [3]. Coordinated expression of P1-2A with controlled 3C^pro^-mediated processing enables efficient formation of genome-free virus-like particles (VLPs) that preserve native antigenic structure while avoiding the requirement to propagate infectious FMDV [4].

Vaccination is central to FMD control and can support regional elimination when applied at scale [5, 6]. However, conventional chemically inactivated FMDV vaccines remain poorly suited to endemic settings: immunity is often short-lived, especially in young cattle; cross-protection among serotypes is limited; antigen integrity is compromised by thermal instability and acid sensitivity, requiring strict cold-chain logistics; and production depends on large-scale propagation of live FMDV in costly, high-containment (BSL-3) facilities [7]. These limitations contribute to suboptimal coverage and recurrent outbreaks, highlighting the need for safe, scalable vaccine platforms better suited to resource-limited settings.

VLPs are a leading next-generation FMD immunogens, and arguably the immunogen of choice, because they preserve the full repertoire of conformational neutralizing epitopes while remaining non-infectious [8]. Multiple VLP production strategies have been described, including recombinant viral vectors and non-viral heterologous expression systems [9]. However, progress toward a practical cattle vaccine has been limited by platform-specific constraints, including restricted cargo capacity in some vectors, the need for high doses and repeated administration, and inconsistent antigen expression and particle yield [9]. As a result, VLP approaches have not yet delivered a scalable, cattle-suitable platform for high-coverage use in endemic settings.

Bovine papular stomatitis virus (BPSV), a parapoxvirus ubiquitous in cattle and associated with absent or mild, self-limiting disease, is a promising viral vector platform [10]. Its favourable attributes include strict host-species restriction, weak and transient anti-vector immunity, persistence in the host, efficient horizontal transmission with high reinfection potential, and substantial foreign-gene cargo capacity [10-15]. Together, these features support repeat dosing, the possibility of “self-boosting” immunity, BSL-2 manufacture and Differentiating Infected from Vaccinated Animals (DIVA)-compatible vaccination strategies suited to FMD control in resource-limited endemic settings.

Here, we evaluated a recombinant BPSV (rBPSV) expressing FMDV virus-like particles (VLPs). We show that BPSV supports high-level VLP production *in vitro* and induces robust neutralizing antibody responses in BPSV-seropositive calves, supporting further development of BPSV as a cattle-adapted VLP vaccine platform for FMD and other high-impact diseases.

## Methods

### Cells and viruses

OFTu cells were maintained in MEM supplemented with 10% fetal bovine serum and standard antibiotics at 37°C with 5% CO_2_. Wild-type bovine papular stomatitis virus (BPSV; C5 strain) [10] and the recombinant BPSV described below were propagated and titrated on primary ovine fetal turbinate (OFTu) cells, with titers calculated by the Reed and Muench method and expressed as TCID_50_/mL.

### Construction of recombinant BPSV expressing FMDV VLPs

To generate rBPSV, the FMDV A24 Cruzeiro capsid precursor P1-2A (VP4, VP2, VP3, VP1 and 2A) and the viral 3C^pro^ were expressed as a single open reading frame (P1-2A-3C^pro^) and inserted into the BPSV024 locus, an NF-κB inhibitory gene implicated in limiting p65 nuclear translocation [10], by double-crossover homologous recombination between parental BPSV C5 strain and a transfer vector following infection/transfection of OFTu cells (Fig. 1A). Because wild-type 3C^pro^ can mediate off-target cleavage of host proteins (including Histone H3, SAM68, NEMO, eIF4aI and eIF4GI) during P1 processing, thereby compromising cell viability and reducing capsid yield, we used an L127P mutant (3C^proL127P^) which has been reported to reduce 3C^pro^-associated cytotoxicity while retaining efficient processing of P1-2A from multiple FMDV serotypes [4]. The recombination cassette comprised the BPSV024 left and right flanking regions, the vaccinia virus vv7.5 early/late promoter driving the P1-2A-3C^proL127P^ expression cassette, and the vv13.5 promoter driving an enhanced green fluorescent protein (EGFP) reporter. Recombinants were isolated through multiple rounds of limited dilution and plaque-purification (Fig. 1B), and their genome integrity confirmed by PCR and sequencing.

**Figure 1.**
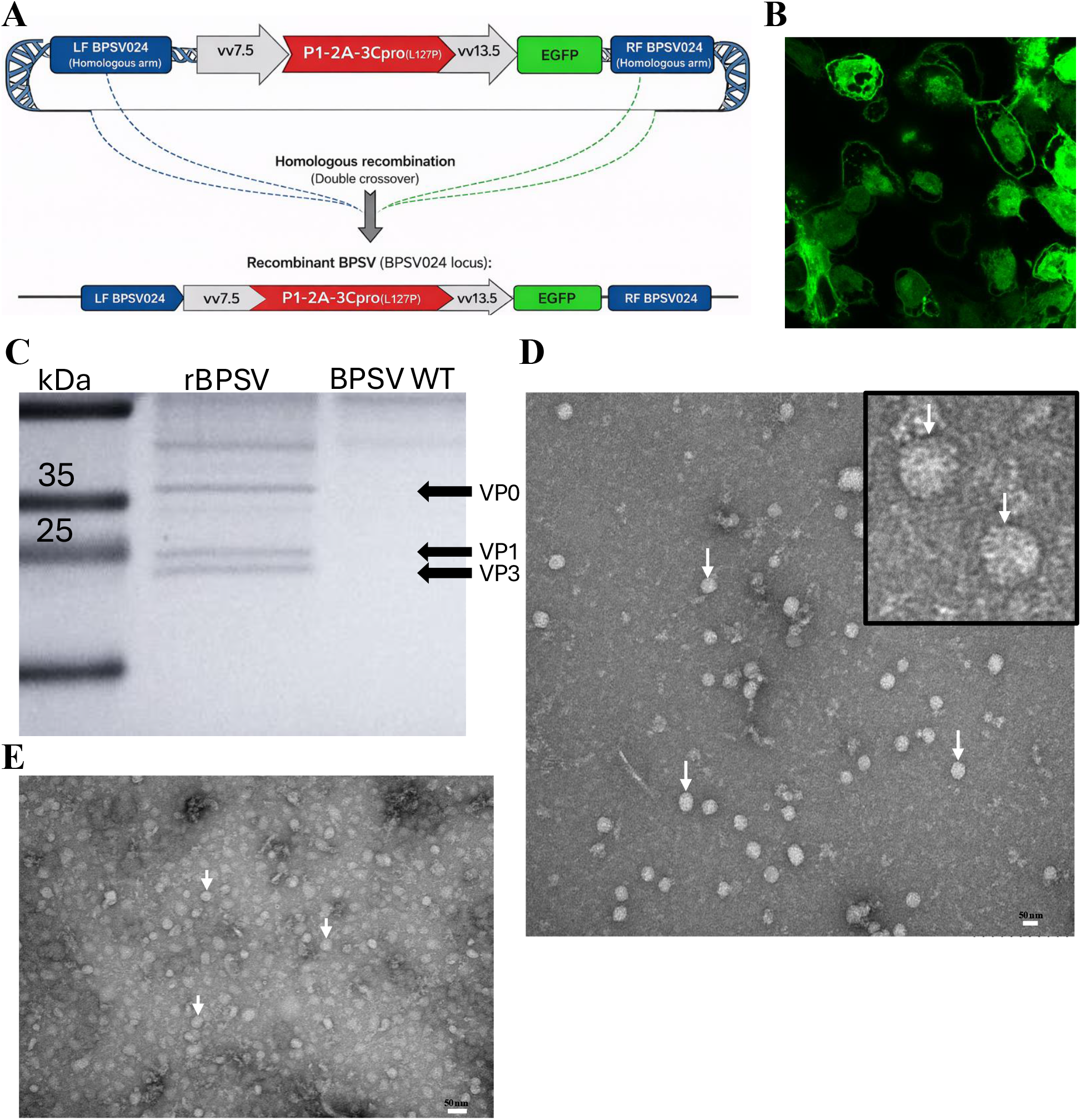
Construction and characterization of recombinant of BPSV expressing FMDV A24 Cruzeiro structural proteins and mutant (L127P) 3C protease [4] (rBPSV-FMDV^P1-2A/3CproL127P^). (A) Schematic of the recombination cassette inserted into the BPSV024 locus, comprising BPSV024 flanking regions, the vv7.5 early/late promoter driving FMDV A24 Cruzeiro P1-2A^3CproL127P^, and the vv13.5 promoter driving EGFP. (B) Fluorescence micrograph of rBPSV-FMDV^P1-2A/3CproL127P^ -infected cells showing EGFP expression. (C) SDS-PAGE of purified VLP preparations from OFTu cells infected with rBPSV-FMDV^P1-2A/3CproL127P^ or wild-type (WT) BPSV, showing VP0, VP1 and VP3 (arrows). (D, E) TEM of negative-stained VLP preparations from infected OFTu cells harvested at 24 hpi (D) and 72 hpi (E), showing abundant ∼25-30 nm icosahedral particles consistent with FMDV empty capsids. Inset in D, higher magnification. Scale bars, 50 nm.

### VLP production, purification and characterization

OFTu cells were infected with rBPSV-FMDV^P1-2A/3CproL127P^ at a multiplicity of infection (MOI) of 0.1 to produce FMDV VLPs. Cells were harvested at 24 and 72 h post-infection (hpi) and lysed by three freeze-thaw cycles. Lysates were clarified by low-speed centrifugation (4,000 *x g*, 20 min, 4 °C) and the supernatants were passed through a 0.45 µm and 0.22 µm membrane filter. VLPs were concentrated by polyethylene glycol precipitation (7% PEG 6000) and purified by ultracentrifugation through a 20% (w/v) sucrose cushion (100,000 *x g*, 4 h, 4 °C). Pelleted VLPs were resuspended in Tris-NaCl-EDTA (TNE) buffer and analyzed by SDS-PAGE with Coomassie Brilliant Blue staining. Particle morphology and size were assessed by transmission electron microscopy (TEM) following negative staining with 1% (w/v) uranyl acetate.

### Immunogenicity of rBPSV-FMDV^P1-2A/3CproL127P^ in cattle

Two calves (4-5-month-old) shown to be BPSV-seropositive by indirect in-house ELISA were vaccinated intramuscularly with 10^7^ TCID_50_ of rBPSV-FMDV^P1-2A/3CproL127P^ on days 0, 21 and 35. Sera collected on days 0, 7, 14, 21, 28, 35, 42, 49 and 56 were analyzed for FMDV-specific humoral responses. Antibodies targeting an immunodominant neutralizing site were quantified by indirect ELISA using synthetic peptides corresponding to the VP1 G-H loop (residues 135-160). Total antibody reactivity to intact capsids was measured by liquid-phase blocking ELISA (LPBE). Functional neutralizing activity was assessed by a conventional virus microneutralization test (VNT) using cell-culture-adapted FMDV strains on BHK-21 cells, as described previously [16]. Neutralizing antibody titers were expressed as the log_10_ reciprocal serum dilution neutralizing 50% of the virus inoculum, calculated using the Reed and Müench method.

To define antigenic breadth, neutralization was evaluated against a panel of viruses spanning increasing antigenic distance, including the homologous strain A24 Cruzeiro, an intra-serotypic heterologous strain (A/Argentina/2001), and a cross-serotypic heterologous strain (O_1_ Campos). Serological data were interpreted in accordance with WOAH guidelines [2] and laboratory-established protective thresholds defined at SENASA (WOAH Reference Laboratory, Argentina) [17, 18]. For A24/Cruzeiro and A/Argentina/2001, the expected percentage of protection (EPP) values were derived from previously established correlations between LPBE titres [17, 19] or VNT titres [20, 21] measured in vaccinated cattle at 60 days post-vaccination and challenge outcomes assessed at 90 days post-vaccination using the Protection against Podal Generalization (PPG) assay [2]. These correlations were generated from groups of 16 vaccinated cattle challenged with the homologous virus. For both total and neutralizing FMDV-specific antibodies, an EPP of ≥75% was used as the reference threshold for antibody titres associated with population-level protection against homologous challenge.

## Results

### Generation of rBPSV-FMDV^P1-2A/L127P3Cpro^ and in vitro VLP production

A stable rBPSV-FMDV^P1-2A/3CproL127P^ was generated with no apparent reduction in viral replication kinetics compared with parental BPSV C5 strain under the conditions tested (data not shown). Infection of OFTu cells resulted in robust expression of VP0 (VP2+VP4), VP1, and VP3 on SDS-PAGE (Fig. 1C). Consistent with proteolytic processing and capsid assembly, TEM revealed abundant 25-30 nm icosahedral particles at 24 hours post-infection (hpi), with marked increase at 72 hpi (Fig. 1D, E). Together, these data indicate efficient *in vitro* assembly of FMDV VLPs.

### Humoral immunogenicity of rBPSV-FMDV^P1-2A/3CproL127P^ in BPSV-seropositive calves

Immunization of two BPSV-seropositive calves with rBPSV-FMDV^P1-2A/3CproL127P^ induced strong FMDV-specific humoral responses (Fig. 2A-C). Consistent with previous observations, no BPSV-associated lesions developed in either calf [10]. Antibodies to the VP1 G-H loop increased steadily after priming, peaked by day 28, and remained elevated through day 56 (Fig. 2A). Total anti-capsid reactivity measured by LPBE increased after the first boost and remained above the defined threshold associated with protection (≥1.9 log_10_) [17, 19] throughout the study period (Fig. 2B). Similarly, functional neutralizing antibody titres against homologous FMDV A24 Cruzeiro exceeded the defined protective threshold (≥1.36 log_10_) [20, 21] following the first boost (Fig. 2C).

**Figure 2.**
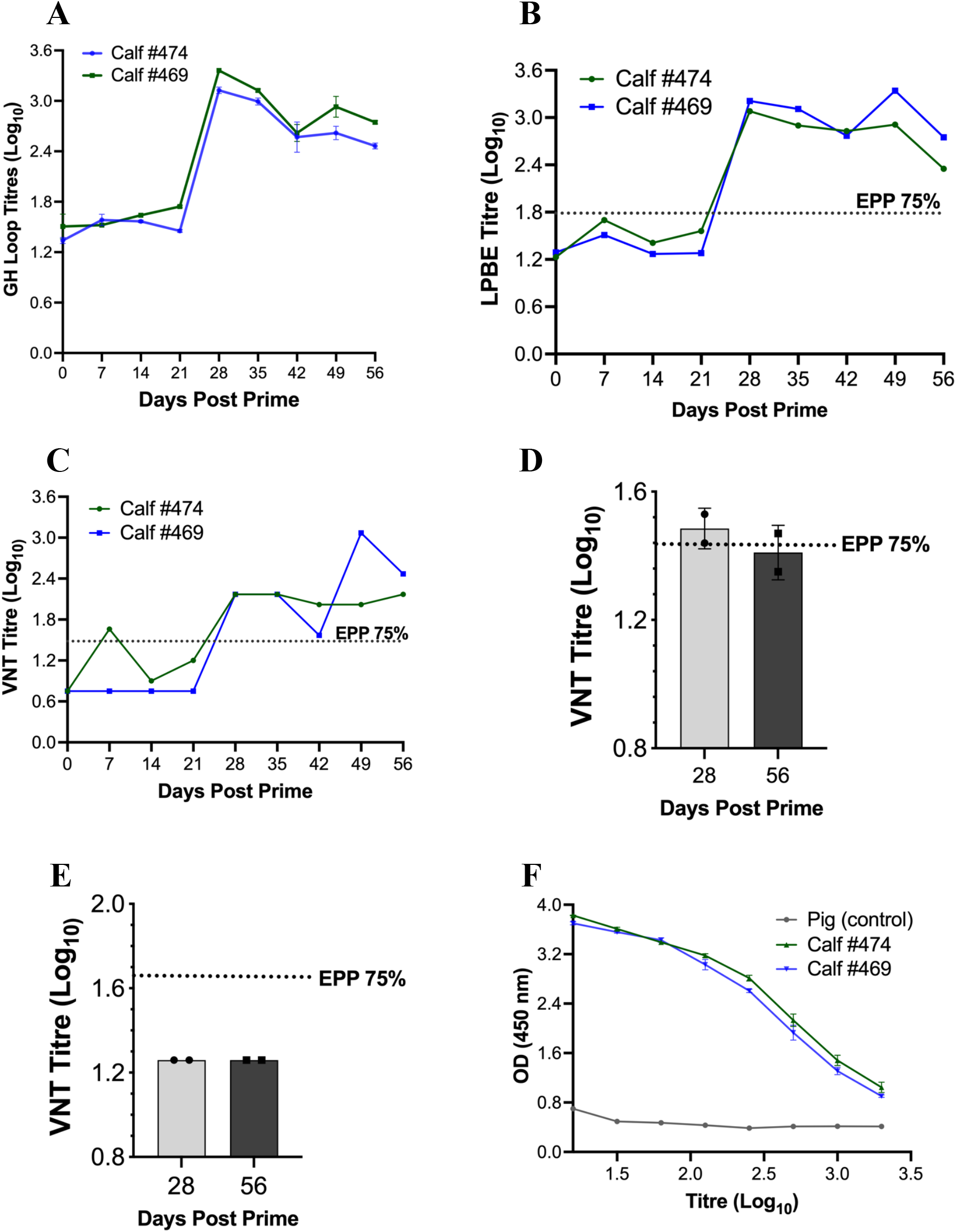
Humoral responses of rBPSV-FMDV^P1-2A/L127P3Cpro^ in BPSV seropositive calves. Two BPSV-seropositive calves were immunized intramuscularly with 10^7^ TCID_50_ of rBPSV-FMDV^P1-2A/3CproL127P^ on days 0, 21 and 35. Sera collected on days 0-56 were analyzed for FMDV-specific antibody responses. (A) VP1 G-H loop peptide ELISA titers (log_10_). (B)Total anti-capsid antibody titers measured by liquid-phase blocking ELISA (LPBE; log_10_); dashed line indicates the 75% expected percentage of protection (EPP). (C-E) Neutralizing antibody titers (VNT; log_10_) against homologous A24 Cruzeiro (C) and heterologous strains A/Argentina/2001 (D) and O_1_ Campos (E). Dashed lines indicate the 75% expected percentage of protection (EPP) for A/24/Cruzeiro (VNT >1.36), A/Argentina/2001 (VNT >1.43) and O_1_ Campos (VNT >1.65). (F) BPSV-specific antibody titers in calves from days 0-56.

Assessment of antigenic breadth demonstrated measurable cross-neutralization of the heterologous intra-serotypic strain A/Argentina/2001, with titers approaching 75% EPP threshold at day 28 before declining toward borderline levels by day 56 (Fig. 2D). In contrast, neutralization of the heterologous cross-serotypic strain O_1_ Campos was not detected at any time point (Fig. 2E), consistent with serotype-specific neutralization. Importantly, pre-existing anti-BPSV antibodies did not prevent induction of FMDV-specific neutralizing responses following vaccination. Collectively, these data show that rBPSV-FMDV^P1-2A/3CproL127P^ induces robust, functional humoral immunity even in animals with prior exposure to BPSV and rBPSV (Fig. 2F).

## Discussion

Conventional inactivated FMD vaccines remain central to control programs but high cost, poor thermostability, short-lived immunity, and limited heterologous protection [7, 9], underscore the need for alternative platforms. This study provides first report that a cattle-adapted parapoxvirus can be engineered to generate high-yield, FMDV VLPs capable of inducing functional immunity in cattle, even in the presence of baseline vector immunity.

The performance of rBPSV-FMDV^P1-2A/3CproL127P^ at the level of antigen production is particularly compelling. BPSV accommodated the ∼4 kb P1-2A-3C^proL127P^ cassette at the BPSV024 locus without an apparent reduction in replication kinetics under the conditions tested, consistent with prior report demonstrating stable foreign-gene insertion in the BPSV genome [10]. Robust VP0, VP1 and VP3 expression together with abundant 25-30 nm icosahedral particles by TEM is consistent with efficient P1 processing and high-level assembly of VLPs in infected cells (Fig. 1C-E). BPSV provides a cattle-adapted platform for *in vivo* VLP production and delivery, avoiding *ex vivo* VLP manufacture and non-bovine-adapted vectors.

Importantly, rBPSV-FMDV^P1-2A/3CproL127P^ induced functional neutralizing antibody titers against homologous A24 Cruzeiro that exceeded the defined protective thresholds (>1.36 log_10_), with LPBE titers persistently above protective threshold (≥1.9 log_10_) (Fig. 2A-C). Based on established serological correlates [17, 19-21], these responses are consistent with protection against homologous challenge in the PPG model. Antigenic breadth analyses showed intra-serotypic cross-neutralization against A/Argentina/2001 (Fig. 2D), with titres approaching the 75% EPP threshold at peak responses, whereas no neutralization was detected against the heterologous serotype O_1_ Campos (Fig. 2E), consistent with the serotype specificity of FMDV neutralizing antibodies [2] . Thus, rBPSV-FMDV^P1-2A/L127P3Cpro^ elicited both strong homologous neutralization and measurable within-serotype cross-neutralization. Vector and cassette optimization, including molecular adjuvants and modulation of viral immunomodulatory genes, may improve vaccine potency and durability of neutralizing responses.

Although anti-vector immunity can limit boosting and repeat immunization [22], BPSV is ubiquitous in cattle and reinfection is common, consistent with weak and transient anti-vector immunity [10, 15, 23]. Accordingly, pre-existing anti-BPSV antibodies did not prevent induction of FMDV-specific neutralizing responses in vaccinated calves (Fig. 2F). Together with manufacture under BSL-2 conditions, compatibility with lyophilization and capacity for large foreign-gene insertion, features above support BPSV as a scalable platform for repeated, multivalent vaccination in endemic settings and for broader veterinary vaccine applications.

Collectively, these data establish proof-of-concept for rBPSV as a cattle-adapted FMDV vaccine platform that combines efficient VLP assembly with neutralizing antibody responses that meet defined protective thresholds [17, 19-21] despite pre-existing vector immunity. Importantly, the persistence, reinfection potential and contact transmission efficiency of BPSV [10, 15, 23] raise the possibility of “self-boosting” immunity.

## Acknowledgement

We thank Dr Manuel Borca for his help, the UNL animal facility staff for technical support, and Bara Altartouri from the UNL Microscopy Core Facility for electron microscopy technical support.

## Author contributions

Conceptualization, H.M.C., G.D., D.L.R., methodology, H.M.C., G.D., B.J.S., S.K., D.M.P-F., D.L.R., investigation, H.M.C., G.D., M.C.M., S.G-N., A.T., B.J.S., D.M.P-F., D.L.R., Data curation, H.M.C., M.C.M., S.G-N., A.T., D.M.P-F., G.D., D.L.R., Funding acquisition, D.L.R., supervision, G.D., D.L.R., All authors participated in revising and approving the final version of the manuscript.

## Ethical approval

The animal study was approved by University of Nebraska-Lincoln Institutional Animal Care and Use Committee (IACUC; protocol #2369). The study was conducted in accordance with the local legislation and institutional requirements.

## Data availability statement

The authors confirm that the data supporting the findings of this study are available within the article.

